# Personalized *Clostridioides difficile* engraftment risk prediction and probiotic therapy assessment in the human gut

**DOI:** 10.1101/2023.04.28.538771

**Authors:** Alex Carr, Nitin S. Baliga, Christian Diener, Sean M. Gibbons

**Affiliations:** Institute for Systems Biology, Seattle, WA, USA; Molecular Engineering Program, University of Washington, Seattle, WA, USA; Departments of Biology and Microbiology, University of Washington, Seattle, WA, USA; Lawrence Berkeley National Lab, Berkeley, CA, USA; Diagnostic and Research Institute of Hygiene, Microbiology and Environmental Medicine, Medical University of Graz, Graz, Austria; Departments of Bioengineering and Genome Sciences, University of Washington, Seattle, WA, USA; eScience Institute, University of Washington, Seattle, WA, USA

## Abstract

*Clostridioides difficile* colonizes up to 30-40% of community-dwelling adults without causing disease. *C. difficile* infections (CDIs) are the leading cause of antibiotic-associated diarrhea in the U.S. and typically develop in individuals following disruption of the gut microbiota due to antibiotic or chemotherapy treatments. Further treatment of CDI with antibiotics is not always effective and can lead to antibiotic resistance and recurrent infections (rCDI). The most effective treatment for rCDI is the reestablishment of an intact microbiota via fecal microbiota transplants (FMTs). However, the success of FMTs has been difficult to generalize because the microbial interactions that prevent engraftment and facilitate the successful clearance of *C. difficile* are still only partially understood. Here we show how microbial community-scale metabolic models (MCMMs) accurately predicted known instances of *C. difficile* colonization susceptibility or resistance *in vitro* and *in vivo*. MCMMs provide detailed mechanistic insights into the ecological interactions that govern *C. difficile* engraftment, like cross-feeding or competition involving metabolites like succinate, trehalose, and ornithine, which differ from person to person. Indeed, three distinct *C. difficile* metabolic niches emerge from our MCMMs, two associated with positive growth rates and one that represents non-growth, which are consistently observed across 15,204 individuals from five independent cohorts. Finally, we show how MCMMs can predict personalized engraftment and *C. difficile* growth suppression for a probiotic cocktail (VE303) designed to replace FMTs for the treatment rCDI. Overall, this powerful modeling approach predicts personalized *C. difficile* engraftment risk and can be leveraged to assess probiotic treatment efficacy. MCMMs could be extended to understand the mechanistic underpinnings of personalized engraftment of other opportunistic bacterial pathogens, beneficial probiotic organisms, or more complex microbial consortia.

## Introduction

The human gut microbiome plays important roles in shaping host metabolism, in the development of chronic diseases, and in preventing opportunistic pathogen colonization and infection ^1–3^. The metabolic versatility of gut bacteria allows for the stable coexistence of hundreds of commensal species within the gastrointestinal tract ^4^. Some species extract energy and nutrition directly from indigestible dietary substrates, like plant fibers or recalcitrant proteins, while others subsist largely on host-derived mucosal glycans or on the vast array of metabolic byproducts produced by primary fiber, protein, and mucus degraders ^5,6^. Saturation of these metabolic niches by commensal microbes can prevent colonization and engraftment by external microbes that may share a similar niche, including pathobionts ^7–9^.

Perturbations to the gut microbiome (e.g., antibiotic use or diarrhoeal events) provide a window of opportunity for pathobiont colonization ^10^, which could in turn lead to the development of disease following subsequent perturbations ^11,12^. One such pathobiont, *Clostridioides difficile*, is the most common hospital acquired gastrointestinal infection in the U.S. ^13,14^. *C. difficile* colonizes as much as 30-40% of community-dwelling adults without causing disease, lying in wait until the opportunity for infection arises ^15,16^. During active *C. difficile* infection (CDI), antibiotic treatment can be effective in suppressing *C. difficile* growth, but antibiotics also disrupt the ecology of the commensal microbiota and potentiate reinfection if *C. difficile* is not completely cleared by the treatment ^11,12^. Thus, an intact gut microbiota that prevents *C. difficile* colonization and engraftment is critical to the host’s defense against CDIs ^3^. This understanding has led to the widespread use of fecal microbiota transplants (FMTs) as a means of combating cases of recurrent CDI (rCDI), where antibiotic treatment proves insufficient ^17^. While the biology of *C. difficile* has been fairly well-characterized in the context of disease, the pre-disease mechanisms of *C. difficile* colonization and engraftment are still poorly understood, as are the factors that govern *C. difficile* decolonization and FMT efficacy ^10^.

There are currently no mechanistically-grounded, generalizable approaches to accurately predicting the engraftment of an exogenous bacterial taxon in the context of a given microbiota. Previous work has leveraged machine learning (ML) to predict the engraftment of FMT donor strains in FMT recipients ^18^. While effective and relatively accurate, this kind of quasi-black-box ML approach does not provide a means of understanding the molecular mechanisms that facilitate or prevent engraftment. Here, we present an alternative approach to engraftment prediction that leverages microbial community-scale metabolic models (MCMMs), which provide detailed, mechanistic information on the ecological interactions within individual microbiota that prevent or facilitate engraftment, in addition to generating accurate engraftment predictions.

Genome-scale metabolic models and classical flux balance analysis (FBA) have been invaluable tools for exploring how environmental conditions impact the metabolic capacities of individual bacterial taxa grown *in vitro* ^19^. However, extending these methods to complex, multi-species communities has proved to be a challenge. Recently, we developed an approach called cooperative tradeoff flux balance analysis (ctFBA), which leverages microbiome compositional and dietary constraints to rapidly estimate steady-state community-scale metabolic fluxes ^20,21^. Here, we use publicly available 16S amplicon and shotgun metagenomic data from *in vitro* and *in vivo* studies with both known and unknown *C. difficile* colonization dynamics, along with our community-scale metabolic modeling framework, called MICOM ^20^, to build and test MCMMs to estimate *C. difficile* engraftment potential within a given microbiome and dietary/environmental context. We present novel insights into how *C. difficile* can occupy three discrete metabolic niches across commensal communities, what metabolic interactions within gut bacterial communities promote or prevent colonization, and we show how we can predict potential responders and non-responders to a defined probiotic cocktail that has recently shown efficacy in the treatment of rCDI ^22^. Overall, MCMMs provide a novel path towards predicting *C. difficile* engraftment risk. Furthermore, these models can be leveraged to design precision dietary or probiotic interventions aimed at decolonizing individuals who are already carrying *C. difficile* and preventing engraftment in those who are not. Finally, we suggest that MCMMs could enable precision engineering of the gut microbiome through personalized engraftment predictions for other pathobionts beyond *C. difficile*, probiotic bacterial strains, or for entire microbial consortia (e.g., FMTs from different donors), in the context of a specific diet.

## Results

### Development of an in silico invasion assay to simulate C. difficile colonization

To simulate the colonization of *C. difficile* we developed an *in silico* invasion assay that leverages microbiome relative abundance data, manually curated genome-scale metabolic models of gut bacteria from the AGORA database, constraints on the diet/environment, and the MICOM modeling framework (Fig. 1A) ^20,23^. Here, we used several existing 16S amplicon and shotgun metagenomic sequencing data sets to validate our approach ^10,18,24–27^. Amplicon sequencing data is often limited to genus-level resolution in the taxonomic classifications of amplicon sequence variants (ASVs). Even with metagenomic sequencing data, species level resolution can be poor when mapping to available species-level metabolic models. Generally, only the most abundant or prevalent commensal bacterial species are well represented. Therefore, we focused on genus-level MCMMs, to approximate the community metabolic context, for our invasion assays (see Methods) ^28^. However, we also evaluated the use of species-level MCMMs using *in vitro* 16S amplicon and stool shotgun metagenomic data (Fig. S1A). Briefly, strain-level metabolic models from AGORA were combined at the genus or species level, to account for potential coexistence of multiple strains from a given genus or species within an individual and to reduce potential bias from arbitrarily selecting individual strain models. Using this approach ∼75% of reads, on average, could be mapped to a genus-level metabolic model within the AGORA database from 16S amplicon data, while ∼90% of reads, on average, could be mapped using shotgun metagenomic data. At the species level, only ∼50% of reads could be mapped, on average, using the same *in vivo* metagenomics data, while 100% of reads could be mapped for *in vitro* communities profiles using 16S amplicon sequencing (i.e., because type strains were used in these experiments), illustrating the inherent limitation of using species level models for *in vivo* applications and further justifying our focus on genus-level analyses (Fig. S1A). To simulate the invasion of *C. difficile* into these model communities, a pan-species model of *Clostridioides*, representing all four common *C. difficile* strains present in the AGORA database (including hypervirulent and non-epidemic strains), was introduced at a relative abundance of 10% (see below for justification for this percentage), while other community relative abundances were decreased proportionally to approximate a minor perturbation in community-wide biomass (Figure 1A). Growth simulations were then performed using a medium representing an average European diet (i.e., a standard developed-world diet appropriate to the cohorts studied here), with fluxes of metabolites known to be absorbed in the small intestine decreased by 90%, as previously described for *in vivo* samples ^20^. To represent the anaerobic basal broth (ABB) used to culture the *in vitro* communities, a curated medium was constructed using a previously described representation of Luria-Bertani (LB) broth as a template and the known composition of ABB (see Methods for further details) ^27,29^. Growth rates were estimated using ctFBA, as implemented in MICOM, which uses a regularization step and allows for a suboptimal community growth rate in order to achieve a more realistic growth rate distribution across the community ^20,21^. Import and export fluxes were estimated using parsimonious enzyme usage FBA (pFBA) ^20^.

**Figure 1.**
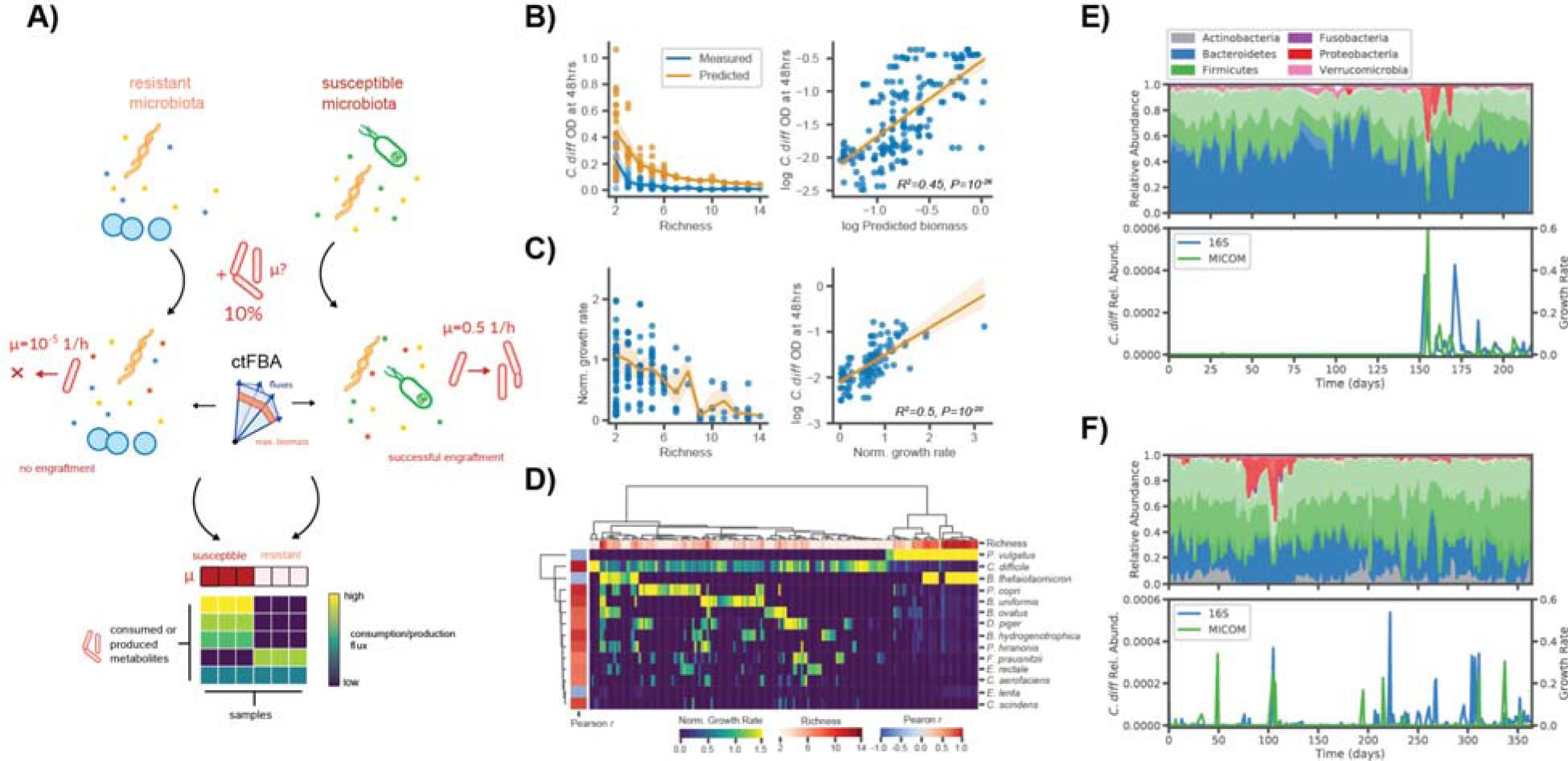
*in silco* invasion assay accurately predicts *C. difficile* engraftment *in vitro* and *in vivo*. (**A**) Schematic illustrating the *in silico* invasion assay workflow leveraged in this study. Personalized microbial community-scale metabolic models (MCMMs) are supplemented with 10% of a pan-genus *Clostridioides* model to simulate an invasion event and ctFBA was used to predict *C. difficile* engraftment and metabolic fluxes. (**B**) Measured *C. difficile* biomass at 48 hours from *Hromada et al*. and predicted biomass, computed from MCMM-predicted *C. difficile* growth rate and an exponential growth model, across a gradient of *in vitro* community richness. Mean trend lines and standard deviation are displayed for both measured (blue) and predicted (yellow) biomass. Relationship between log_10_ predicted biomass and log_10_ biomass. Ordinary least squares fit and 95% confidence interval are displayed, as well as regression R^2^ and *p*-value (**C**) Normalized *C. difficile* growth rate along a community richness gradient and the relationship between normalized *C. difficile* growth rate and measured log_10_ biomass. Mean trend line and standard deviation are displayed for the relationship between normalized *C. difficile* growth rate and community richness. Ordinary least squares fit and 95% confidence interval are displayed, as well as regression R^2^ and *p*-value, for the relationship between normalized *C. difficile* growth rate and measured log_10_ biomass. (**D**) Biclustered normalized growth rates for species across conditions. Species normalized growth rate is displayed using blue-to-yellow heatmap coloring for each sample. Sample community richness is displayed using white-to-red heatmap coloring on the top row of the plot. Pearson correlation coefficients between *C. difficile* normalized growth rate and the normalized growth rate of each other species are displayed using blue-to-red heatmap coloring in the leftmost column of the plot. Colorbars at the bottom of the blot indicate the scales for the various measures. (**E**) Donor A time series taken from David *et al.* displaying daily fluctuations in microbiome composition over a period of several months. Composition is displayed, colored by phylum-level annotations (different shading indicates taxonomic families). At day 150, Donor A experienced a diarrheal event and was subsequently colonized by *C. difficile*. Estimates of *C. difficile* relative abundance from 16S sequencing and MICOM-predicted *C. difficile* growth rates are displayed. (**F**) Time series from Donor B from the same study, who was apparently colonized by *C. difficile* (at very low relative abundances, near the limit of detection) throughout the sampling period.

Personalized MCMMs were constructed for each sample and the potential for *C. difficile* engraftment was quantified as the model-inferred growth rate. ctFBA has a single free parameter that needs to be chosen, the tradeoff between community-wide growth rates and individual, taxon-specific growth rates. Assuming that most taxa detected at appreciable abundances in a gut microbiome are actively growing *in vivo* and *in vitro*, a trade-off value was selected by choosing the minimal deviation from optimal community growth for which >70% of genera obtained non-zero growth rates on average (Fig. S1B). We found that with a trade-off value of 0.8 (i.e., 80% of maximal community biomass production) resulted in a mean fraction of genera with non-zero growth of >70% using genus-level models constructed from 16S amplicon data. This same non-zero growth fraction was achieved with a trade-off value of 0.9 for the genus-level models constructed from metagenomics data. The 16S and metagenomics specific tradeoff values were also used for the species level models derived from these data types, respectively. At the species-level these values were associated with much higher growth fractions (∼90% and 95% for 16S amplicon and shotgun metagenomics, respectively; Fig. S1B).

We first validated our approach using an *in vitro* 16S amplicon data set where *C. difficile* was co-cultured with communities of human commensal gut bacteria at differing levels of species richness ^27^. We found that MCMM-inferred *C. difficile* biomass (calculated using the MCMM-inferred *C. difficile* growth rates, assuming exponential growth over a fixed interval, see Methods) accurately reflected a negative trend in measured *C. difficile* abundance along a gradient of increasing species richness and was significantly correlated with empirically measured *C. difficile* biomass after 48 hours of growth (Fig. 1B; ordinary least squares (OLS) R^2^=0.45, *p* <10^−16^). Similarly, we found that the normalized *C. difficile* growth rate (normalized to the overall MCMM-inferred community growth rate, see Methods) decreased as a function of community richness and was a even stronger predictor of the empirically-observed *C. difficile* biomass after 48 hours of growth (Fig. 1C; ordinary least squares (OLS) R^2^=0.5, *p* <10^−16^). Biclustering of the inferred species growth rates suggested that growth of *C. difficile* was largely suppressed by *B. thetaiotaomicron, P. vulgatus*, and *E. lenta in vitro*, as indicated by the strong negative associations between the growth rates of these species and the growth rate of *C. difficile* (Pearson *r* <-0.45, p <0.01 for species). Positive associations between the growth rates of *C. difficile, C. scindens*, and *P. hiranonis* (Pearson *r* >0.8, p <10^−6^ for all species) were inline with the observation that these species tend to share overlapping metabolic niches. The previously-observed suppressive effects of *C. scindens* and *P. hiranonis* on *C. difficile* were not apparent within our modeling framework ^27^, likely due to the fact that pH and its effects on growth are not easily captured by FBA models. However, while *in vitro* acidification is a strong environmental force in anaerobic batch culture, we do not expect variation in pH to be as prominent in the buffered environment of the human colon.

To further validate our approach *in vivo* we leveraged a time series with a known *C. difficile* colonization event (Fig. 1E) ^10,30^. We found that estimated *C. difficile* growth rates were at or below the limit of solver accuracy (<10^−6^, which effectively indicates an absence of growth) in samples collected prior to colonization and comparable to growth rates of other dominant genera in samples taken after the initial colonization event (Welch’s t-test *t*=-3.29, *p*=0.003, for comparison of log_10_ growth rates before and after the known colonization event; Fig. 1E). Furthermore, we saw patchy engraftment predictions in a second individual that was known to be colonized by *C. difficile* at a low level (i.e., near the limit of detection) throughout a time series (Fig. 1F). We also assessed the importance of propagule pressure ^31^ (i.e., the relative abundance at which the invasive taxon is introduced into the models) and found that below 10% relative abundance, agreement between growth rate estimates and measured abundances was poor (Fig. S1C). Thus, propagule pressure plays an important role in predicted engraftment success ^27^.

Based on these initial results (Fig. S1), we decided to use a fixed tradeoff value of 0.8 and a *C. difficile* invasion fraction of 10% for all subsequent analyses of 16S amplicon data (i.e., the highest value at which >70% of the community showed a positive growth rate). Using the same heuristic, a tradeoff value of 0.9 was chosen for the metagenomics analyses, along with a 10% *C. difficile* invasion fraction (Fig. S1B).

### in silico invasion assay accurately predicts C. difficile colonization potential in rCDI patients pre- and post-FMT

We applied our *in silico* invasion model to two separate datasets (16S amplicon and metagenomic shotgun sequencing, respectively) of rCDI patients who received FMTs and were subsequently followed over time ^18,26^. These data provided an additional test of MCMM performance and a means to explore the metabolic features associated with community-scale colonization susceptibility or resistance across a larger population. Given that all individuals in the rCDI cohorts had experienced multiple recurrent infections, we expected pre-FMT microbiomes from these patients to be more susceptible to invasion. Additionally, both of the original studies found that nearly all the patient microbiomes transitioned to a compositional state that was much closer to the healthy controls post-FMT than to their pre-FMT compositional states ^18,26^. Thus, we expected post-FMT samples would be less susceptible to invasion. For the sake of comparison of results across studies, genus-level MCMMs were used for both analyses. Across both data sets, rCDI patients had significantly higher MCMM-predicted *C. difficile* growth rates prior to FMT treatment than they did post FMT treatment (Fig. 2A; Welch’s t-test *t*=-3.19, *p*=0.001 for comparison of pre-FMT vs. post-FMT). Predicted *C. difficile* growth rates were negatively associated with Shannon diversity, albeit weakly (Fig. 2B; OLS R^2^=0.05, *p*=0.01), which is in line with prior empirical observations indicating that lower diversity communities are more susceptible to *C. difficile* colonization and the development of rCDI ^9,27,32,33^.

**Figure 2.**
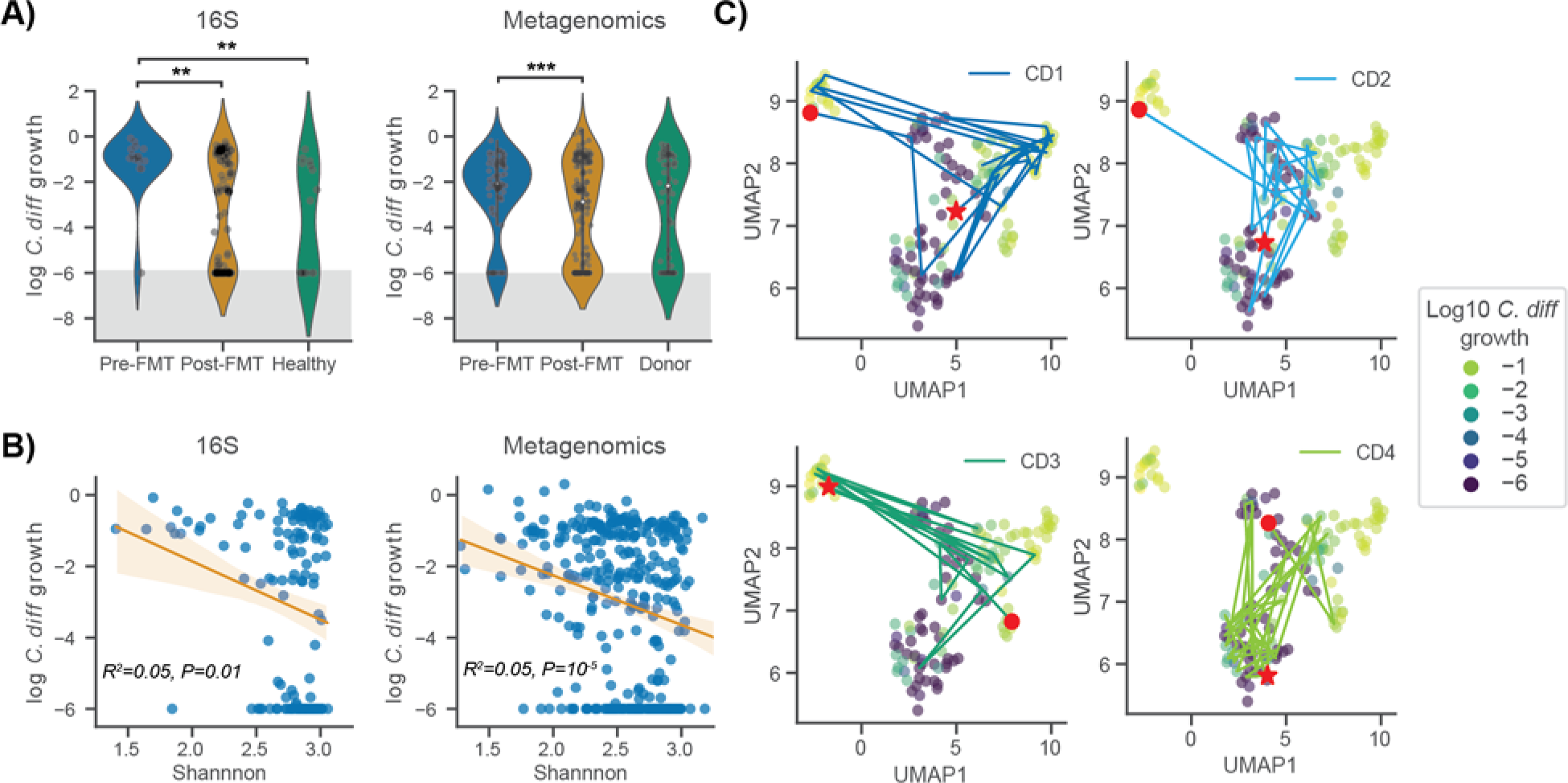
*C. difficile* growth rate predictions capture importance of community context for patient recovery from CDI. (**A**) Violin plots displaying predicted *C. difficile* log_10_ growth rate distributions across patient disease status categories. Data from Weingarden *et al.* and Ianiro *et al.* were leveraged to validate the approach using both 16S amplicon and metagenomic sequencing. Gray shading indicates the numerical accuracy of the solver (values below 10^−6^ can not be distinguished from zero and are considered negligible). Bars indicate comparisons for which differences were significant using Welch’s t-test. *, *p* <0.05; **, *p* <0.01; ***, *p* <0.001. (**B**) Relationship between predicted *C. difficile* log_10_ growth rate and Shannon diversity across datasets. Ordinary least squares fit and 95% confidence interval are displayed, as well as regression R^2^ and *p*-value. (**C**) Two-dimensional representation of community import fluxes prior to *in silico* invasion using UMAP colored by log_10_ growth rates of *C. difficile* following *in silico* invasion. Patient trajectories are displayed, each with a red circle representing the patient’s starting point (prior to FMT), and a red star representing the patient’s end point (post recovery).

The community-scale import flux profile prior to *in silico* invasion was predictive of *C. difficile* growth rate following invasion in Weingarden *et al.* (Fig. 2C). High-dimensional community-scale import flux profiles were projected into a two-dimensional space using the Uniform Manifold Approximation and Projection (UMAP) technique (Fig. 2C) ^34^. The UMAP projection provides a visual means of identifying patterns in the high dimensional import flux space. The closer points are to one another in this ordination the more similar their import flux profiles are. Thus, clusters of points in the UMAP can represent distinct metabolic environments across samples. The ordination plot indicated that *C. difficile* appears to grow well in more than one metabolic environment, when colonizing different individuals. Indeed, we saw that the predicted metabolic environments occupied by *C. difficile* could vary within an individual over time (Fig. 2C). For most patients in the Weingarden *et al.* cohort, there was a transition from colonization-susceptibility pre-FMT to colonization-resistance post-FMT (Fig. 2A,C). We next examined the different metabolic niches that *C. difficile* was able to exploit when colonizing individuals across both rCDI-FMT cohorts, to better understand this phenotypic plasticity.

### C. difficile is predicted to occupy three distinct metabolic niches within the human gut microbiome

To characterize *C. difficile* colonization-associated niches and identify the potential for multiple metabolic strategies associated with its growth, we examined *C. difficile* import fluxes with high variance (log_10_ flux variance >=4.5) across the rCDI-FMT cohorts. Using this criteria the high variance metabolites identified across the 16S and metagenomic data sets were highly consistent (Fig. 3). We identified 31 and 44 high-variance metabolite fluxes from the 16S and metagenomics analyses, respectively. Of these, 27 were shared, 4 were unique to the Weingarden *et al.* (16S) analysis, and 17 were unique to Iraniro *et al.* (metagenomics) analysis. Biclustering of the high variance import flux data and an examination of how the apparent clusters associated with growth rates revealed that *C. difficile* makes use of multiple metabolic strategies (Fig. 3). Three major clusters were observed across patient samples and cohorts. We designate these three clusters as “high growth”, “moderate growth” and “no growth” (Fig. 3). The high growth cluster in both cohorts included many of the pre-FMT samples and was characterized by consistently high import fluxes for all the metabolites identified as most strongly coupled to *C. difficile* growth across all models. The moderate growth cluster showed a sparser metabolite consumption profile. For example, ornithine and fructose were rapidly consumed in the high growth cluster, but showed almost no consumption in the moderate growth cluster (Fig. 3). As expected, very few metabolites were consumed by *C. difficile* above the zero-threshold of 10^−6^ in the no growth cluster (Fig. 3).

**Figure 3.**
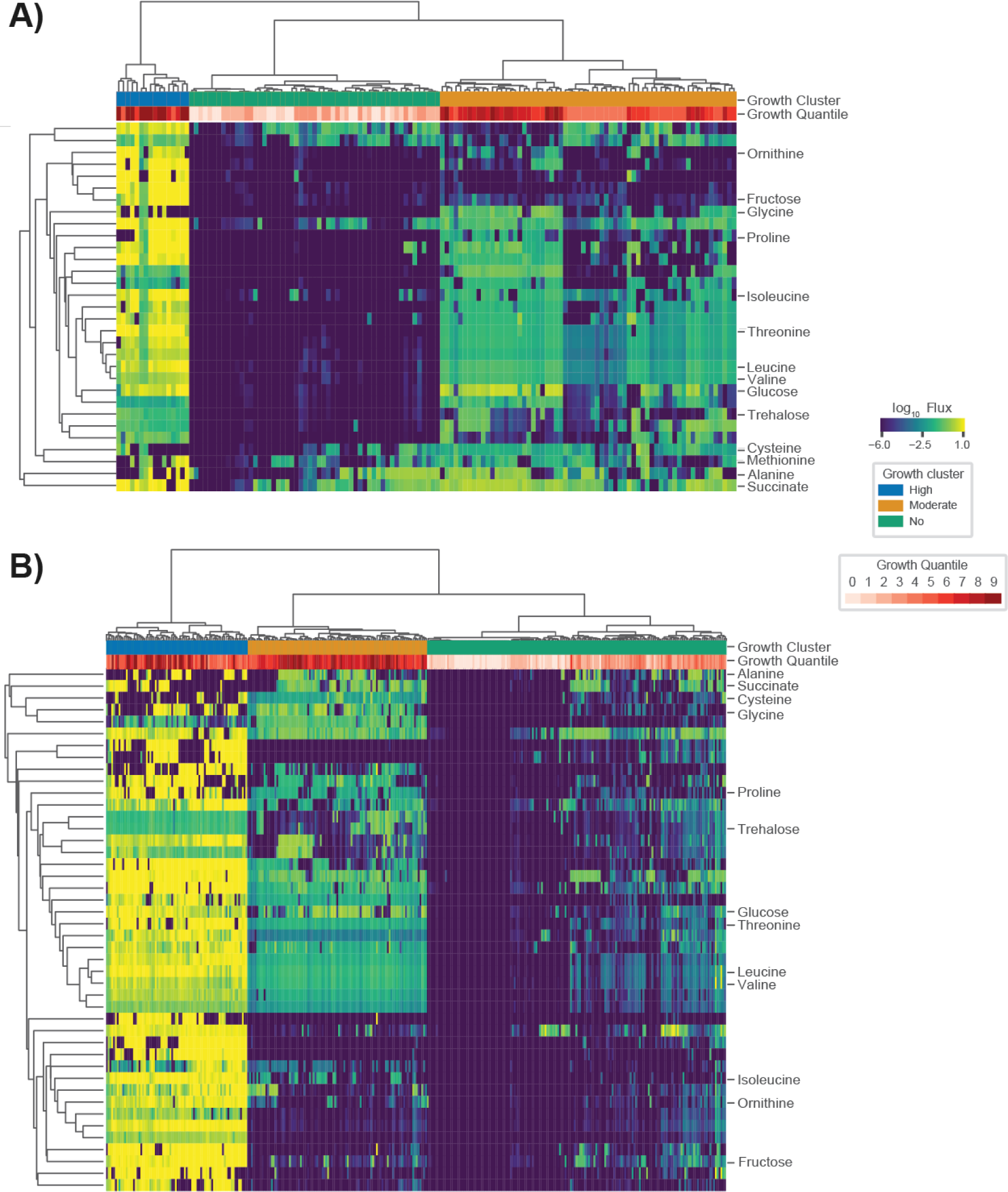
*C. difficile* occupies multiple metabolic niches across communities. (**A**) Biclustered *C. difficile* log_10_ import fluxes from Weingarden *et al.*, where each row is the import flux of a particular metabolite and each column is a patient sample. (**B**) Biclustered *C. difficile* log_10_ import fluxes from Ianiro *et al.*, where each row is the import flux of a particular metabolite and each column is a patient sample. In each heatmap, imports for which the log variance across samples was >=4.5 are displayed. Fluxes across samples are displayed using blue-to-yellow heatmap coloring. *C. difficile* log_10_ growth rate quantiles are displayed in white-to-red heatmap coloring for each patient sample in the top row of each plot. Additionally, three coarse grain growth clusters are noted. These growth clusters represent “high growth”, “moderate growth”, and “no growth” phenotypes.

The metabolic strategies employed by *C. difficile* within the MCMMs showed convergence with several observations from the literature. For example, we found that metabolites known to promote growth of *C. difficile in vivo* (e.g., succinate, ornithine, and trehalose) were preferentially utilized when available and were associated with higher pathobiont growth rates ^35–37^. In addition, the consumption of amino acids valine, leucine, glycine, glutamate, glutamine, and proline were associated with higher *C. difficile* growth rates in the MCMMs, implying that *C. difficile* employs Strickland fermentation in one of its growth modes, which has been observed empirically ^38^.

Following up on these findings we examined how cooperative and competitive interactions within MCMMs contributed to *C. difficile* colonization. To accomplish this, we examined the import and export fluxes of metabolites associated with *C. difficile* colonization (e.g., amino acids, ornithine, succinate, etc.). Genera that produced metabolites consumed by *C. difficile* likely promote its growth, while those consuming *C. difficile* growth-associated metabolites may be in direct competition. For ornithine and succinate, we found that cooperative and competitive interactions are context-dependent, varying across samples. The genus *Phocaeicola*, for instance, produces ornithine in some samples, which is in turn consumed by *C. difficile*, while in other contexts it consumes ornithine, competing with *C. difficile* (Fig. 4A). Meanwhile, *Roseburia*, and *Faecalibacterium* compete with *C. difficile* for ornithine, but these genera also produce succinate and leucine in some contexts, which *C. difficile* consumes (Fig. 4A-B). Thus, the overall community context, rather than the presence or absence of any single taxon, appears to be the most important factor in determining the metabolic strategies used by *C. difficile* and can lead to emergent competitive or cooperative interactions, which can either hinder or promote colonization.

**Figure 4.**
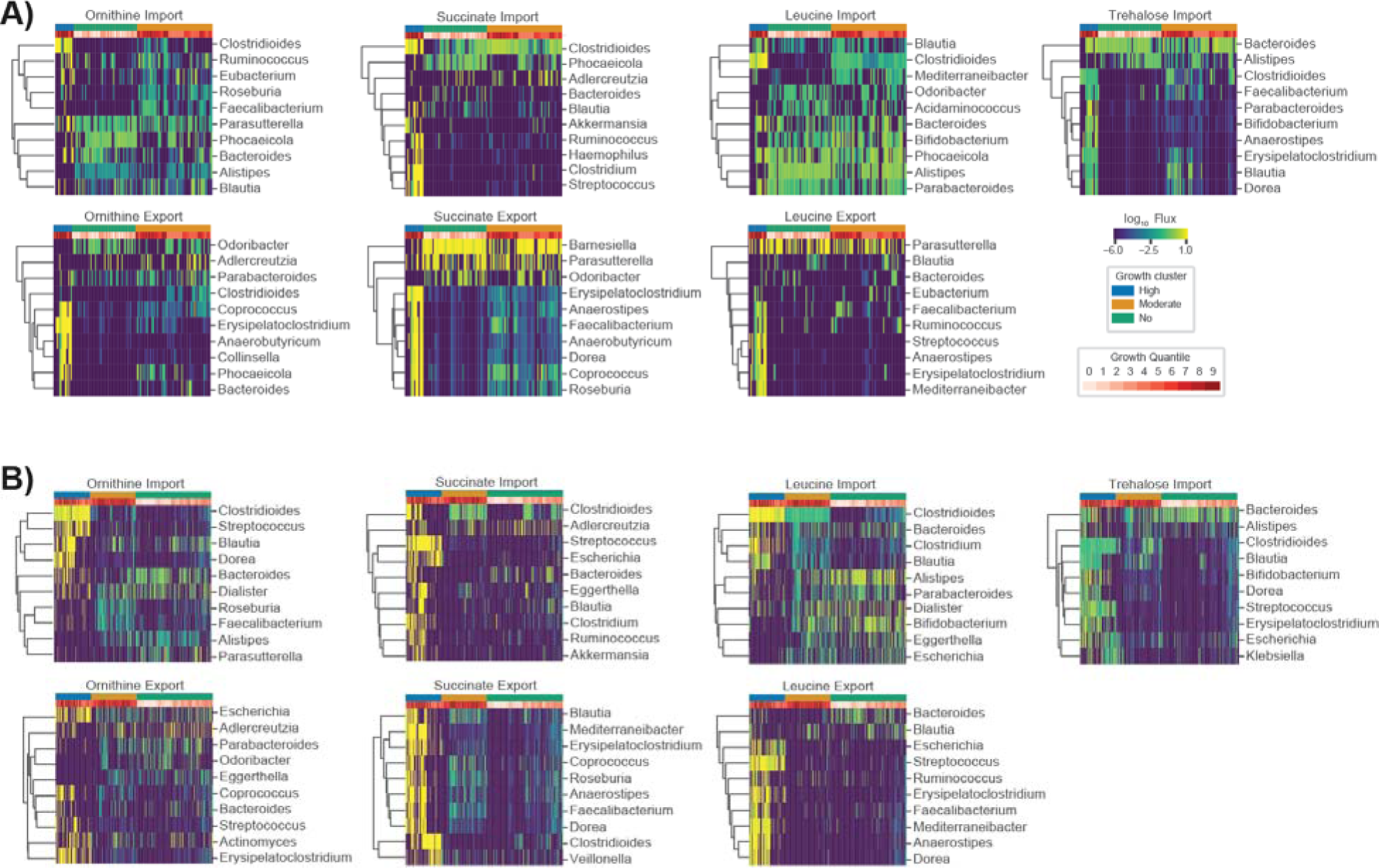
Import and export fluxes of key metabolites across communities highlight the context-dependency of key *C. difficile* competitors and cooperators. (**A**) Biclustered community import and export fluxes of specific metabolites associated with *C. difficile* colonization from Weingarden *et al.*, where each row is a genus and each column is a patient sample. (**B**) Biclustered community import and export fluxes of specific metabolites associated with *C. difficile* colonization from Ianiro *et al.*, where each row is a genus and each column is a patient sample. For each heatmap the top 10 genera with the highest mean import or export flux across samples are displayed. Fluxes across samples are displayed using blue-to-yellow heatmap coloring. *C. difficile* log_10_ growth rate quantiles are displayed in white-to-red heatmap coloring for each patient sample in the top row of each plot. Additionally, three coarse grain growth clusters are noted. These growth clusters represent “high growth”, “moderate growth”, and “no growth” phenotypes.

Finally, we assessed whether or not compositional variation in the microbiome alone could explain observed differences in MCMM-predicted *C. difficile* growth rates. We found that compositional variation was a modest predictor of estimated *C. difficile* growth rate (out-of-sample least absolute shrinkage and selection operator (LASSO) regression R^2^=0.37 for the Weingarden *et al.* rCDI-FMT cohort, see Methods). Meanwhile, the import flux derived growth clusters (e.g., “high growth”, “medium growth”, and “no growth” groups) could explain the vast majority of the variance in predicted *C. difficile* growth rates (analysis of variance (ANOVA) R^2^=0.94 for the CDI-FMT cohort). However, this comparison between model-estimated growth rates and growth clusters is a bit circular (i.e., both are derived from the model), so as mentioned above, we also observed that MCMM-estimated *C. difficile* growth rates were able to explain 50% of the variance in empirical *C. difficile* biomass measurements *in vitro* (Fig. 1C). In summary, these results suggest that compositional information alone is not sufficient for consistently accurate engraftment predictions.

### Associations with C. difficile growth provide insights into the role of community context

In order to assess the consistency of the *C. difficile* growth clusters, we leveraged four independent *in vivo* 16S amplicon data sets, including the time series and the Weingarden *et al.* rCDI-FMT studies presented above (Figs. 1-4), along with two large cross-sectional cohorts (i.e., the American Gut and Arivale cohorts), covering a total of 14,862 individuals ^24,30,39,40^. We evaluated growth and flux predictions generated across all four data sets and found that *C. difficile* fell into the same three clusters that were identified in the rCDI-FMT data sets, representing no growth, moderate growth, and high growth (Fig. 5A).

**Figure 5.**
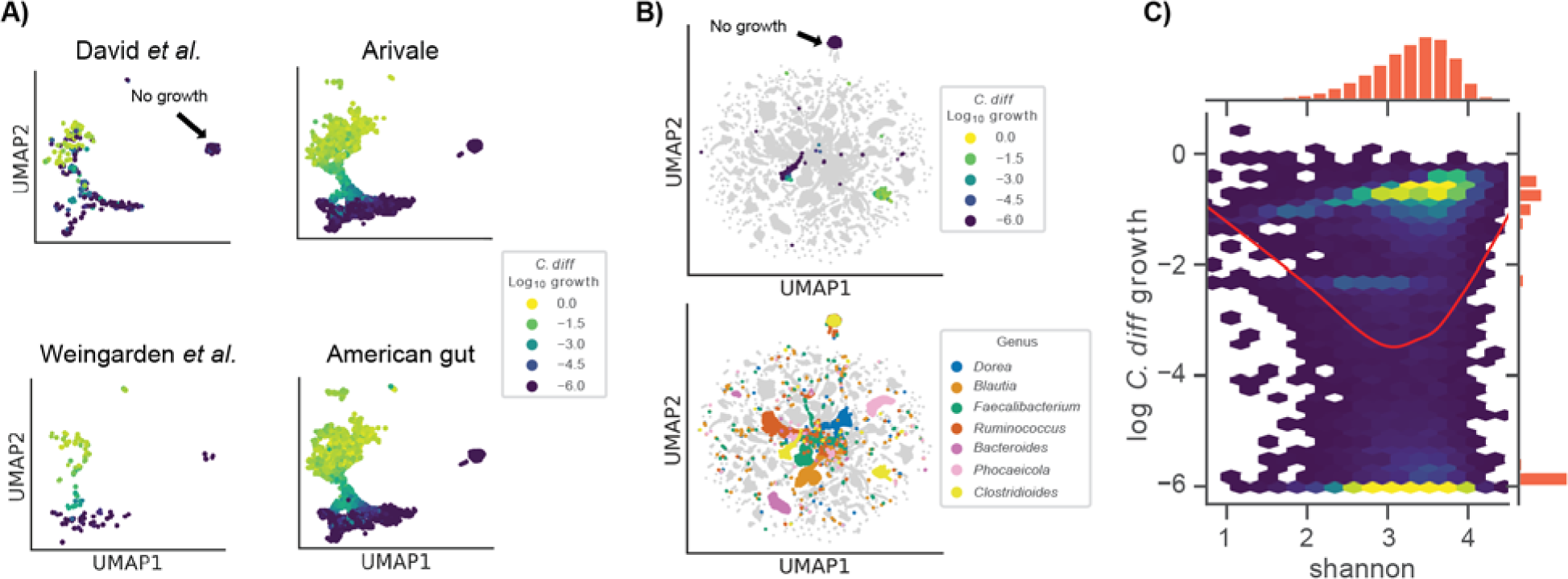
Growth niches across large four independent cohorts challenged *in silico* with *C. difficile*. (**A**) Two dimensional representation of log_10_ *C. difficile* import fluxes using UMAP across four independent data sets. Colors denote *C. difficile* growth rate ranging from low (blue) to high (yellow). The position of the no growth cluster is indicated. (**B**) Two dimensional representation of log_10_ genus import fluxes using UMAP across all datasets. Top panel displays log_10_ *C. difficile* growth rate within the context of all other genera. Bottom panel colors *C. difficile* along with six genera of interest: *Dorea*, *Blautia*, *Faecalibacterium*, *Ruminococcus*, *Bacteroides*, and *Phocaeicola*. The position of the no growth cluster is indicated. (**C**) Two-dimensional hexagonal binning of log_10_ *C. difficile* growth rate and community alpha diversity (Shannon index). Red trend line indicates a LOWESS fit to the log_10_ *C. difficile* growth rate and community Shannon diversity data.

To further contextualize the metabolic niche of *C. difficile*, we integrated model outputs for all four 16S data sets. Specifically, we looked at import fluxes across all genera. Most genera formed unique clusters in the UMAP projection, suggesting that each genus had a single metabolic niche that was consistent across datasets (Fig. 5B). Within this community context, we found that *C. difficile* still fell into three very distinct clusters (Fig. 5B). Several genera that showed some of the strongest competitive and cooperative interactions with *C. difficile* across data sets, *Blautia*, *Faecalibacterium*, *Ruminococcus* and *Dorea*, clustered near one another in import flux space, with some overlap, indicating that these commensal taxa shared a similar metabolic niche (Fig. 5B). Meanwhile, *Bacteroides* and *Phocaeicola*, two apparent *C. difficile* competitors, formed clusters largely separate from those formed by the other genera. Several of these taxa also occupied niche space near the *C. difficile* moderate growth cluster, supporting the potential for competitive interactions between *C. difficile* and these genera. The *C. difficile* high growth cluster was more separated from other commensal genera, suggesting that this growth mode is more specialized in the context of the surrounding community (Fig. 5B).

We next explored gut community diversity and predicted *C. difficile* growth rates across the four data sets. Specifically, we looked at Shannon diversity, which integrates species richness and evenness and is commonly used to quantify gut microbiome alpha-diversity. Lower Shannon diversity is commonly associated with disease states, like diarrhea, while higher diversity has generally been associated with diverse plant-based diets and overall better health ^32^. However, constipated individuals generally have higher gut microbiome alpha-diversity as well, suggesting that there may be an optimal range of alpha-diversity across healthy individuals ^39,41^. Our initial analysis using the rCDI-FMT cohort suggested a monotonically negative relationship between predicted *C. difficile* growth rate and Shannon (Fig. 2B). However, the integrated data sets, which spanned a wider range of diversity, showed a U-shaped relationship between Shannon diversity and predicted *C. difficile* growth rate (Fig. 5C). Intermediate levels of Shannon diversity were associated with the lowest predicted growth rates, on average, with higher average growth at the upper and lower tails of the distribution (Fig. 5C). The relationship between Shannon diversity and predicted growth rate suggests extremes in either direction on the diversity scale are, on average, more permissive to *C. difficile* engraftment.

### Blood metabolites and clinical labs associated with MCMM-predicted C. difficile colonization susceptibility

We next sought to identify potential blood-based markers that were significantly associated with MCMM-predicted *C. difficile* growth rates. Previous work has shown that circulating blood metabolites can be leveraged to predict gut microbiome alpha-diversity ^39^. We identified several blood metabolites and clinical chemistries significantly associated with MCMM-predicted *C. difficile* growth rates in the Arivale cohort, after adjusting for common covariates (i.e., sex, age, and BMI) and correcting for multiple tests (FDR *q* <0.05). These included two secondary bile acids, an unannotated metabolite previously associated with the abundance of the family *Eggerthellacea*, and several red blood cell-associated clinical chemistries (Fig. S1D)^42^. Unfortunately, while significant, these blood-based markers, along with sex, age, and BMI, collectively accounted for only ∼5% of the variance in MCMM-predicted *C. difficile* growth rates. Thus, it appears MCMM-based estimates of *C. difficile* engraftment, constrained by fecal microbiome data, cannot be readily replaced with commonly measured clinical chemistries or blood metabolites.

### MCMMs predicts engraftment heterogeneity of probiotic cocktail designed to treat rCDI

As a final proof-of-concept for our modeling framework we simulated a probiotic intervention using a previously validated probiotic cocktail designed to treat rCDI ^22,43^. The probiotic, referred to as VE303, was composed of 8 commensal Clostridia strains and shown to be effective at treating CDI in mice ^43^. This probiotic was also shown to be safe, well-tolerated, and effective in reducing rCDI incidence in humans ^22^. An earlier study in both mice and humans found that engraftment of VE303 strains was optimal following antibiotic pretreatment ^43^. With these facts in hand, we designed a simulated intervention that mimicked the treatment found to be most effective by Dsouza *et al* ^43^. We were only able to identify metabolic models for 6 of the 8 strains in VE303 in the AGORA database ^23^. We leveraged the Weingarden *et al.* rCDI-FMT dataset to test this six-member probiotic cocktail, paired with *in silico* invasion by *C. difficile*. The probiotic cocktail was introduced to patient samples, alongside 10% *C. difficile*, at a total relative abundance of 50%, which was evenly distributed among the six strains. We also simulated vancomycin treatment by reducing the abundance of *C. difficile* and all commensal genera known to be impacted by vancomycin treatment ^44^ by 90%. We found that a combined probiotic and antibiotic intervention most effectively suppressed the growth of *C. difficile* in both the moderate and high *C. difficile* growth rate clusters (Fig. 6A).

**Figure 6.**
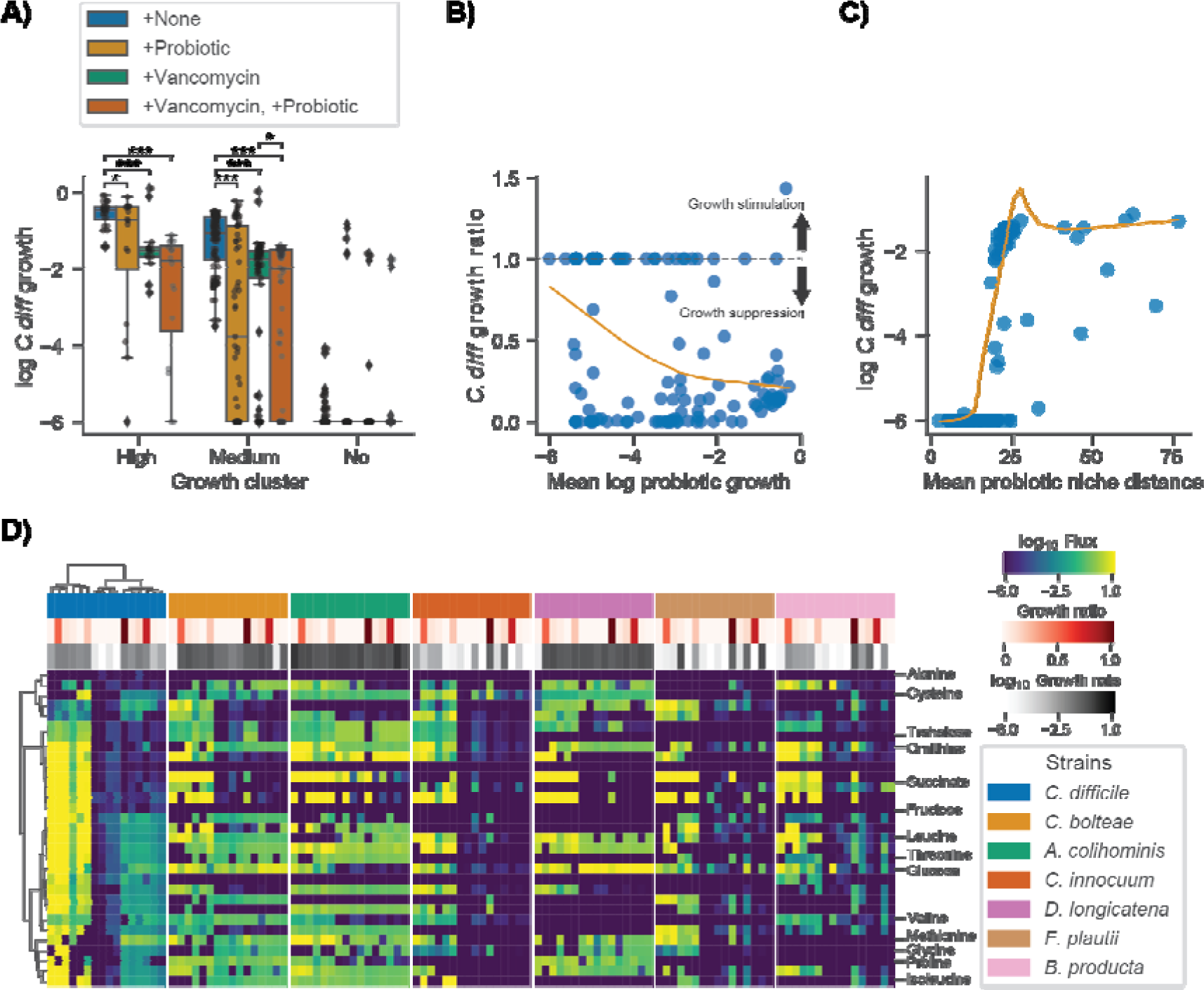
Simulated probiotic intervention effectively suppresses *C. difficile* growth *in silico*. (**A**) Box plots displaying log_10_ *C. difficile* growth rate across growth clusters and simulated interventions. Growth clusters are those identified by bi-clustering of *C. difficile* import fluxes using the Weingarden data (Figure 3). Conditions include +None (no intervention control), +Probiotic (introduction of 6 strain probiotic previously identified as an effective treatment for CDI at a total relative abundance of 50% equally distributed across the strains), +Vancomycin (90% reduction of *C. difficile* relative abundance as well as all genera known to be impacted by Vancomycin), and +Vancomycin, +Probiotic (introduction of 6 strain probiotic in combination with simulated vancomycin treatment). Bars indicate comparisons for which differences were significant using the Wilcoxon signed-rank test. *, *p* <0.05; **, *p* <0.01; ***, *p* <0.001. (**B**) Relationship between *C. difficile* growth ratio and mean log_10_ probiotic growth rate. *C. difficile* growth ratio is the growth rate of samples in the +Vancomycin, +Probiotic intervention relative to +None. Values below 1 indicate growth suppression by the probiotic and values above 1 indicate growth stimulation. The dashed line marks the value at which no effect is observed (1). Orange trend line indicates a LOWESS fit to the *C. difficile* growth ratio and mean log_10_ probiotic growth rate. (**C**) Relationship between log_10_ *C. difficile* growth rate and mean probiotic niche distance. Niche distance was calculated using the euclidean distance of log_10_ import flux vectors of each probiotic strain relative to *C. difficile* on a per sample basis. Orange trend line indicates a LOWESS fit to the log_10_ *C. difficile* growth rate and mean probiotic niche distance. (**D**) Biclustered log_10_ import fluxes for *C. difficile* and probiotic strains for samples previously identified as “high growth”, where each row is the import flux of a particular metabolite and each column is a patient sample. Imports displayed are those previously identified as important for *C. difficile*. Color bars indicate sample *C. difficile* growth ratio and strain specific log_10_ growth rate. Ordering of samples and metabolites is the same across heatmaps and based on biclustering of *C. difficile* data.

To better understand the mechanism of action of the probiotic cocktail, we assessed the growth characteristics and the niche proximity of the probiotic strains in relation to *C. difficile.* We found that suppression of *C. difficile* growth occurred more readily when the average growth of the probiotic strains was high (>10^−4^, Fig. 6B) and when the average niche distance between the probiotic strains and *C. difficile* was low (<25 import flux Euclidean distance, Fig. 6C). We also found that, relative to other genera, several of the probiotics strains occupied niches closer to *C. difficile*, although these niche distances could vary widely for each organism depending on their community context (Fig. S2). Finally, we compared the import fluxes of the probiotic strains and *C. difficile* for the metabolites identified as important for *C. difficile* growth (Fig. 3). This analysis showed that, in addition to occupying niches similar to *C. difficile*, several of the probiotic strains directly competed for metabolites important for *C. difficile* growth, such as succinate, ornithine, and trehalose (Fig. 6D). Cumulatively these results suggest that metabolic competition is the mechanism by which the probiotic cocktail suppressed *C. difficile* growth, which is consistent with the emerging consensus in the field ^9,43,45^. Finally, we found that certain probiotic strains were more or less likely to engraft in an individual (Fig. 6D), and that this engraftment/growth was generally associated with *C. difficile* suppression (Fig. 6B), which indicates that MCMMs can be leveraged to identify responders and non-responders prior to these kinds of probiotic interventions.

## Discussion

In this study, we provide a framework for predicting *C. difficile* engraftment risk in the human gut microbiome using MCMMs. While we focus on *C. difficile*, due to its clinical importance, this approach could be extended to other opportunistic bacterial pathogens, probiotic organisms, or even entire communities, in the case of FMTs. We were able to show how our approach predicts expected longitudinal and cross-sectional variation in *C. difficile* colonization potential *in vitro* and *in vivo* (Figs. 1-2), using both shotgun metagenomic and 16S amplicon sequencing data sets, and we provide insights into the metabolic strategies leveraged by *C. difficile* in different ecological contexts (Figs. 3-4). Our analysis not only recapitulates known metabolic associations with *C. difficile* growth (e.g., consumption of trehalose, ornithine, and succinate; Fig. 3), it suggests additional associations (e.g., importance of reduced sulfur compounds like cysteine, Stickland fermentation reactants, and utilization of dietary sugars, like fructose; Fig. 3). Additionally, we show that competition and cooperation with community members can prevent or promote colonization of *C. difficile*, and that many of these associations are highly context-dependent (Fig. 4).

Consistent with the idea that simple metrics of community structure and composition alone are not effective predictors of colonization susceptibility, we found that community compositional variation was a modest predictor of estimated *C. difficile* growth rate and the relationship between alpha diversity and estimated *C. difficile* growth rate was nonlinear (Fig. 5C). Not only did low diversity communities tend to be more invasible, as might be expected due to putative non-saturation of the metabolic niche space ^9^, but high diversity communities were also more prone to *C. difficile* engraftment (Fig. 5C). In high diversity communities, successful colonization may be due to changes in the interaction landscape, like a higher rate of niche-construction in higher-diversity systems, which would be in line with the diversity-begets-diversity hypothesis ^46^. An intermediate range of alpha-diversity seems to be optimal for mitigating *C. difficile* colonization potential (Figure 5C). Overall, these complex mappings between community composition and pathobiont engraftment risk underscore the necessity of systems-scale tools, like MCMMs, that are capable of synthesizing this complexity.

Several genera were found to engage in cooperative and competitive interactions with *C. difficile* across MCMMs. *Blautia*, *Faecalibacterium*, and *Dorea* were all shown to benefit *C. difficile* through production of key metabolites that it consumes, like succinate and leucine, but were also capable of competition for other metabolic resources (Fig. 4). Meanwhile *Rumminococcus*, *Bacteroides* and *Phocaeicola* growth rates were often negatively associated with *C. difficile* engraftment, competing for some of the same metabolites that *C. difficile* consumed at high rates (Fig. 1D, 4). Contextualizing these results through analysis of individual taxon import fluxes across studies we found that *Blautia*, *Faecalibacterium*, *Phocaeicola, Rumminococcus,* and *Dorea* share similar niches with one another, with *Phocaeicola being the most divergent from the others*. In most cases, these niches did not overlap with *C. difficile*, but in a subset of individuals all five occupied niches states overlapping or similar to *C. difficile* (Fig. 5B). Thus, while we observed competition for some key metabolites, on a global scale, the majority of the metabolic niche space used by *C. difficile* tends not to overlap with its apparent competitors (Fig. 5B). These results highlight how flexible commensal gut bacteria are in adapting their import fluxes to the communities in which they reside, which in turn suggests why so many taxa are able to coexist. These results also suggest that colonization resistance is an emergent feature of multi-species metabolic interactions, and not strongly driven by any particular pairwise interaction.

In addition to developing a simulation framework to predict engraftment we sought to identify blood-based clinical chemistries and blood metabolites that were associated with MCMM-inferred *C. difficile* growth rate. We found three blood metabolites that were independently associated with predicted *C. difficile* growth rates. These included two secondary bile acids and an unannotated metabolite. One of the secondary bile acids, isoursodeoxycholate, has previously been positively associated with the abundance of *Bacteroides* ^39^, and was negatively associated with predicted *C. difficile* growth rates. This result is in line with the apparent competition between *Bacteroides* and *C. difficile* in our MCMMs. We also identified several clinical labs negatively associated with predicted growth rate (Fig. S1D). However, together with age, sex, and BMI, these features only accounted for ∼5% of the variance in predicted growth rates. Thus, while these features may be signatures for colonization susceptibility in the blood, their clinical relevance is limited at this time.

Finally, we demonstrated that a probiotic intervention (i.e., 6/8 strains from VE303), which recently showed positive efficacy results in a double-blinded, placebo-controlled clinical trial for the treatment of rCDI ^22^, suppresses the growth of *C. difficile in silico* in most people in the Weingarden *et al.* rCDI-FMT study (Fig. 6). We also showed that the mechanism of action for this particular probiotic is likely competition for metabolites essential for the growth of *C. difficile*, as many of the probiotic strains occupy niches close to *C. difficile* and directly compete for these metabolites, such as succinate and ornithine, in samples where growth suppression was observed (Fig. 6). Furthermore, analysis of niche distances between *C. difficile* and other genera across donors suggest selecting strains from *Blautia and Dorea* (e.g., including *B. producta and D. longicatana,* from VE303), in addition to *Anaerostipes*, *Roseburia*, and *Faecalibacterium*, could be leveraged to design better individual-specific probiotic cocktails capable of suppressing *C. difficile* and rescuing VE303 non-responders (Figs. S2 and 6B). These results illustrate how MCMMs may be powerful tools for assessing the individual-specific efficacy of clinically-relevant probiotics, in addition to understanding personalized pathobiont colonization susceptibility.

Future work should test this MCMM framework in the design of precision interventions to prevent engraftment of *C. difficile* and other pathobionts, to design precision probiotic interventions, and to improve the production of beneficial microbial metabolites, like short-chain-fatty-acids, or reduce the production of undesired metabolites, such as hydrogen sulfide or trimethylamine ^47–49^. In summary, MCMMs present a promising new path forward in engineering the ecological composition and metabolic outputs of microbiota to prevent or treat disease.

## Methods

### External Data Collection and Processing

Data used in this study came from six sources. This included both cross-sectional and time series 16S amplicon and shotgun metagenomics sequencing data from Hromada *et al.*, David *et al.*, Weingarden, A. *et al*., Iraniro *et al.,* the American Gut (McDonald, D. *et al.*), and a former scientific wellness program run by Arivale, Inc.^18,24,26,27,30,40^. Publicly available 16S amplicon sequence data and associated metadata were downloaded from the sequence read archive (SRA). Additionally, de-identified 16S amplicon sequence data, associated metadata, and paired blood-based clinical chemistries and metabolomics were obtained for 2,687 research consenting individuals that were formerly participants in the Arivale wellness program. Raw 16S amplicon sequence data were processed using QIIME2 (v2020.11.1). In brief, the QIIME2 workflow consisted of read demultiplexing using the command *qiime tools import*, and an associated manifest table for each study describing read metadata followed by read quality assessment using *qiime demux summarize*. Read quality assessment was used to determine trimming parameters for subsequent denoising using the QIIME2 implementation of DADA2 via the command, *qiime dada2 denoise-single* or *qiime dada2 denoise-paired*, for single and paired reads respectively. The first 10 bases were trimmed from all reads and reads were truncated to a length where median quality score was >20 (100-150 base pair for the data leveraged). Following denoising, data were reformatted into a table format using the command *qiime metadata tabulate*, and representative sequence taxonomy was inferred using a custom NCBI classifier with the command *qiime feature-classifier classify-sklearn*. The NCBI classifier was trained using 16S 515f-806r V4 regions extracted from all available bacterial NCBI genomes. To train the classifier 515f-806r regions were extracted from NCBI sequences using the command *qiime feature-classifier extract-reads*, followed by the command *qiime feature-classifier fit-classifier-naive-bayes* using the extracted V4 sequences and a table of known taxonomies. Pre-processed data from Iraniro *et al.* were provided by the authors upon request and included sample taxonomic relative abundance inferred using Metaphlan4 and associated sample metadata. See Iraniro *et al.* for further details on their data processing pipelines ^18^. Sample taxonomic relative abundance and associated metadata from Hromada *et al.* were obtained from their supplemental data. See Hromada *et al.* for further details ^27^. For source code and tables of processed data refer to the Github repository listed below in Data and Source Code Availability.

### Model Construction and growth simulations

To construct community level metabolic models sample specific taxonomic abundance profiles inferred from 16S amplicon or shotgun metagenomic sequencing were summarized at the genus or species level and mapped to genus or species level metabolic models from the AGORA database (v1.03) using MICOM (v0.25.1). Taxa with a relative abundance less than 0.1% were omitted from community models. An *in silico* media previously designed to represent an average western diet was applied which defined the bounds for metabolic imports by the model communities for all *in vivo* analyses ^20,23^. For the *in vitro* communities a custom medium was developed to reflect the ABB medium used experimentally. The custom medium was developed using a previously defined template designed for the representation of Luria-Bertani (LB) medium, this contained metabolite concentration and fluxes for both yeast extract and tryptone ^29^. These details were used to convert the quantities of yeast extract and peptone present in the experimental medium anaerobic basal broth (ABB) to fluxes. The additional defined components were included with fluxes equal to the millimolar concentration per hour. Finally, additional components were added to the constructed medium so that all 13 species used in the *in vitro* communities could achieve a growth rate of 10^−3^ or greater using a routine implemented in the MICOM function *complete_db_medium*. Across all conditions, growth rates were inferred using cooperative tradeoff flux balance analysis (ctFBA). In brief, this is a two-step optimization scheme, where the first step finds the largest possible biomass production rate for the full microbial community and the second step infers taxon-specific growth rates and fluxes, while maintaining community growth within a fraction of the theoretical maximum (i.e., the tradeoff parameter), thus balancing individual growth rates and the community-wide growth rate^20^. For all models in the manuscript that leveraged 16S amplicon data we used a tradeoff parameter of 0.8. A tradeoff parameter of 0.9 was used for all the results derived from shotgun metagenomics data. These parameter values were chosen by identifying the largest tradeoff value (corresponding to the smallest deviation from maximal community biomass) that allowed >70% taxa to grow (growth rate >10^−6^). Import and export fluxes were estimated using parsimonious enzyme usage FBA (pFBA) and a defined medium constructed to represent an average European diet or ABB ^20^. pFBA further constrained simulation results by requiring genera to utilize the lowest overall flux through their networks to achieve maximal growth ^19^. For source code and tables of processed data refer to the Github repository listed below in Data and Source Code Availability.

### Biomass estimation and growth rate normalization for in vitro communities

To allow better comparison between model predictions and experimental results two growth rate transformations were employed. These methods were implemented to account for the fact that batch growth (in which cultures grow on a limited quantity of nutrients) is not compatible with the steady state assumption employed by the modeling framework (which assumes continuous supply of nutrients). Estimation of biomass was leveraged to compare growth rates with experimentally measured end point optical densities. This consisted of using a simple exponential growth model of the form *OD_f_ = OD_i_ e^kt^*, where *OD_f_* is the final predicted biomass, *OD_i_* is the initial biomass, *k* is the MCMM inferred growth rate, and *t* is time in hours. The second transformation employed was a growth rate normalization procedure, which accounted for differences in community growth rates across samples. Growth rates were divided by the MCMM inferred community growth rate, which was computed from the sum of relative growth rates of each taxon in the system (e.g., the product of growth rate and relative abundance), 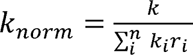.

### Probiotic Intervention

To model probiotic intervention a combination of strains previously shown to be effective at suppressing the growth of *C. difficile* in mice were used ^43^. Metabolic models for six of the eight stains in the VE303 cocktail described by Dsouza *et al.* were identified in the AGORA database and intervention was simulated by introducing them along with *C. difficile* to individual samples. A total probiotic fraction of 50% was used, which was evenly distributed among the six strains. This fraction was determined to be the most effective at suppressing the growth of *C. difficile* growth *in silico* for the samples tested (data not shown). Vancomycin treatment was simulated by reducing the abundance of *C. difficile* and all genera known to be impacted by vancomycin by 90% ^44^. Growth simulations were performed as described above. For source code and tables of processed data refer to the Github repository listed below in Data and Source Code Availability.

### Statistical analysis

Statistical analyses were performed using functions from the python scipy (v1.7.1), seaborn (v0.11.2), sklearn-learn (v0.24.2), umap-learn (v0.5.1), and statsmodels (v0.13.1) packages. Linear associations were performed using the *statsmodels* ordinary least squares function *OLS*, and visualized using the *seaborn* function *regplot*. Least absolute shrinkage and selection operator (LASSO) was performed using the *sklearn Lasso* function and a training-test framework. Data were split into training and test sets (70% of samples were randomly assigned to the training set) and model performance was assessed across a range of regularization values. LASSO training and test set R^2^ values were used to select the model with the best test set R^2^ that did not overfit training data (training R^2^ >> test R^2^). Analysis of variance (ANOVA) was performed using the *statsmodels OLS* and *anova_lm* functions. UMAP dimensionality reduction was performed using the *umap* function from the *umap* package and associated methods with default parameters (i.e., *n_components=2*, *n_neighbors=15*, *metric=’euclidean’*, etc.). Biclustering was performed using the *seaborn* function *clustermap* and the Ward clustering algorithm. Hexagonal binning and associated histograms were generated using the *seaborn* function *jointplot.* Locally weighted scatterplot smoothing (LOWESS) curves were generated using the *lowess* function from *statsmodels* with default parameters. Additional statistical tests included the t-test and Wilcox rank sum test implemented in scipy as *ttest_ind* and *wilcoxon* respectively. For source code and tables of processed data refer to the Github repository listed below in Data and Source Code Availability.

## Data and Source Code Availability

Processed data tables and source code to reproduce the findings presented in this manuscript can be found at https://github.com/Gibbons-Lab/cdiff_invasion. Raw 16S amplicon sequence data from David *et al.*, Weingarden, A. *et al*., the American Gut (McDonald, D. *et al.*) can be downloaded using the sequence read archive (SRA) accession numbers PRJEB6518, PRJEB19996, and PRJEB11419 respectively. Data and metadata from Iraniro *et al.* were provided from the authors upon request. Data from Hromada *et al.* were downloaded from the manuscript supplementary information. Metadata were obtained from manuscript supplementary information, where available. Qualified researchers can access the full Arivale deidentified dataset, including all raw data, supporting the findings in this study for research purposes through signing a Data Use Agreement (DUA). Inquiries to access the data can be made at and will be responded to within 7 business days.

## Acknowledgements

We thank members of the Gibbons Lab for helpful feedback and suggestions on this work. We thank Nicola Segata, Michal Punčochář, and Nicolai Karcher for providing access to their post-processing data files. This work was supported by a Washington Research Foundation Distinguished Investigator Award (to SMG). Research reported in this publication was also supported by the National Institute of Diabetes and Digestive and Kidney Diseases of the National Institutes of Health (NIH) under award no. R01DK133468 (to SMG.).

---

**Figure S1.**
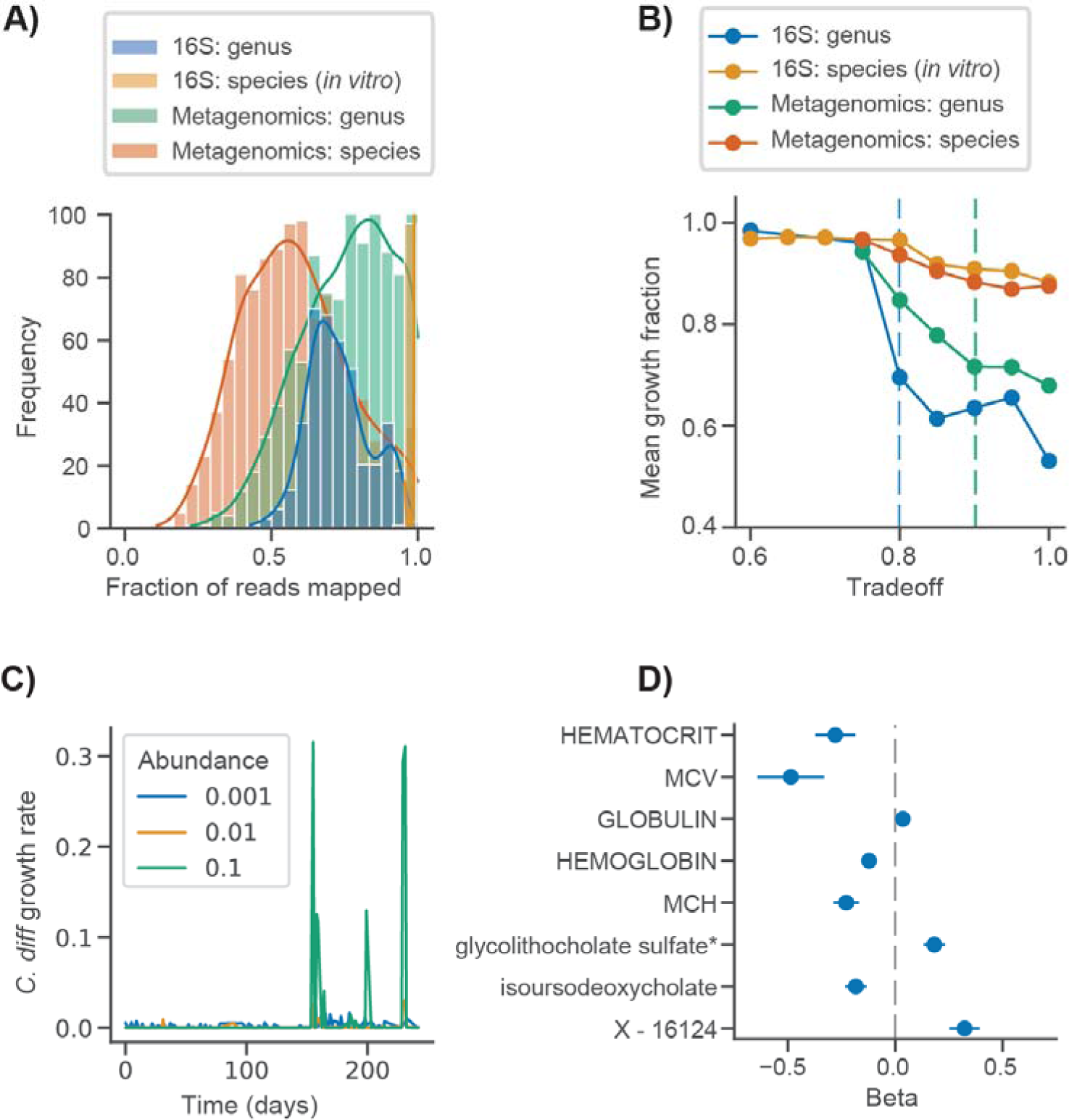
Development of *in silico C. difficile* invasion assay. (**A**) Histograms displaying the fraction of reads mapped at the genus and species levels for the David *et al.* and Hromade *et al.* 16S amplicon data and Ianiro *et al.* metagenomics data using an NCBI reference and metabolic models from the AGORA database. (**B**) Mean growth fraction across samples, datasets, and taxonomic mappings (e.g., fraction of taxa with estimated growth rate >10^−6^) as a function of model tradeoff value. Dashed lines indicated the tradeoff values chosen for subsequent analyses. The blue dashed line indicates the value used for 16S data and green dashed line denotes the value used for metagenomics data. Tradeoff values were chosen such that mean growth fractions of ∼0.7 were achieved using genus level mapping. (**C**) Relationship between *C. difficile* invasion abundance and growth rate for one of the two David *et al.* time series. (**D**) Association coefficients for estimated *C. difficile* log growth rate, blood metabolite concentrations, and clinical labs for the Arivale cohort.

**Figure S2.**
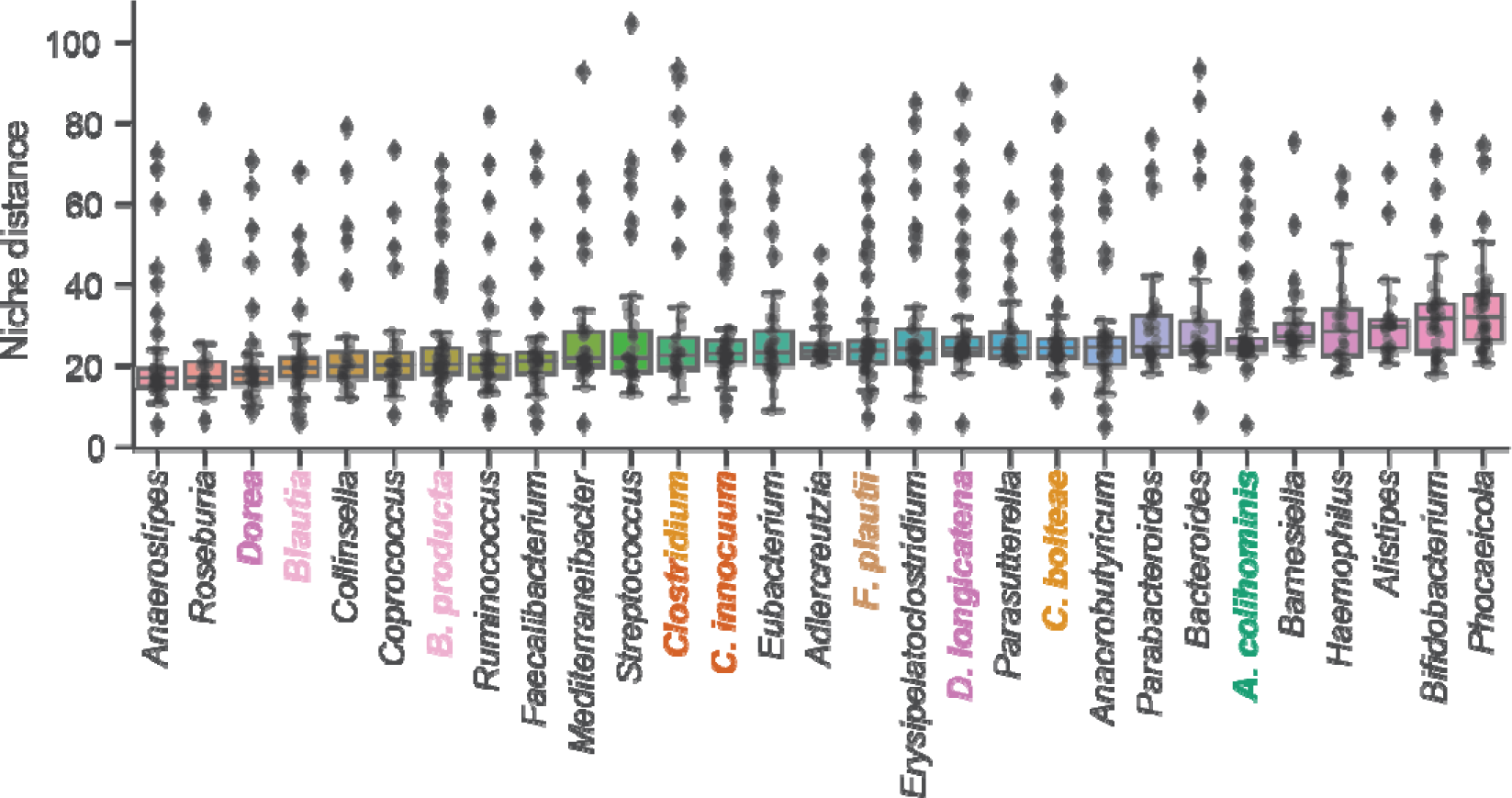
Probiotic strains and associated genera have niche distances close to *C. difficile* relative to unrelated genera. Niche distances of strains and genera, represented as the euclidean distance between flux vectors, relative to *C. difficile* across CDI-FMT cohort samples for which the *C. difficile* growth rate >10^−6^. Genera and strains are ordered by the median niche distance. Probiotic strains and associated genera are colored, consistent with the legend in Fig. 5D. *C. bolteae* and *C. innocuum* are both members of the genus *Clostridium*.

